# A computationally guided approach to improve expression of VHH binders

**DOI:** 10.1101/2024.09.07.611840

**Authors:** Sila Ozdemir, Jessica Tolley, Florian Goncalves, Michelle Gomes, Eli Wagnell, Bruce Branchaud, Viktoriya Dubrovskaya, Sri Ranganathan

## Abstract

The variable heavy chain fragments derived from camelid antibodies, called VHHs or nanobodies, have recently shown promise as high-affinity reagents. They offer higher stability compared to conventional antibodies and fragments thereof. Furthermore, their smaller size (∼15-20 kDa) allows better targeting of molecules localized inside the cell and in crowded environments, like tissues and protein aggregates. Despite these advantages, nanobody clones screened using phage display can suffer from poor soluble expression, which we hypothesized, is due to the presence of hydrophobic hotspots on their surface. In this work, we propose a novel computationally guided workflow for screening and production of nanobody binders for optimized expression. After an initial round of phage display screens against our target (K-Ras), we modeled the lead candidates to generate Spatial Aggregation Propensity (SAP) maps to highlight the hydrophobic hotspots with single amino acid resolution, which were subsequently used to guide mutagenesis of the binders for soluble expression. We followed two approaches to perform point hydrophilic mutations: i) performing point hydrophilic mutations in the hydrophobic hotspots; ii) combining point mutation resulting from a round of random mutagenesis that show favorable SAP scores. Both approaches led a remarkable increase in soluble expression which allowed production and characterization of their binding to their target (K-Ras) on soluble ELISA, and biolayer interferometry. We observed that the latter approach resulted in clones with stronger binding affinity compared to the former approach. Our results emphasize the need to perform a round of random mutagenesis to identify point mutations, which can then be used in an *in-silico* guided pipeline to identify the right combination of mutations for high soluble expression.

## 1. Introduction

Affinity binders are essential reagents and the workhorses of biological research, from academic research to clinical diagnostics and therapeutics. Traditionally, full-size IgG molecules and their fragments have been the go-to affinity binders for assays such as ELISA, Flow Cytometry, Western blot and immunofluorescent staining assays.[1] However, smaller affinity binder scaffolds have been considered for the “next-generation” of affinity reagents for their numerous advantages over conventional antibodies, including smaller size, higher thermodynamic stability, amenability to re-engineering, access to target sites, and ability to penetrate tissues and cells.[2-4] Specifically, single domain antibody binders found in camelids, also known as nanobodies or variable domain of heavy-chain antibodies (VHHs), have recently become popular affinity reagents due to their natural availability, in addition to their above-mentioned advantages.[5, 6]

The quality of any affinity binder, characterized by their specificity, affinity and functional activity, is pivotal for the precision in their application, such as a diagnostic test or achieving their therapeutic potential. Even though VHHs are single heavy chain only fragments containing only three complementary determining regions (CDRs) compared to the six in IgGs, they have been shown to bind their targets with high affinity (K_D_ < nM) and specificity.[7] This has led to rapid development of VHH based affinity binders as potential therapeutics for diseases, including SARS-coV2,[8] Alzheimer’s disease,[9] and cancer.[10]

Our target in this study is K-Ras4B (hereafter referred to as K-Ras), one of the most important oncogenic proteins. K-Ras is one of the four proteins in the small Ras GTPase family,[11] and serves as upstream molecular regulators of very complex downstream kinase pathways such as MAP and PI3K/AKT ensuring cells homeostasis. Since the RAS proteins play an important role in cellular signaling, their dysregulation due to mutations lead to uncontrolled cell proliferation, making it an important target of cancer diagnostics and therapeutics.[12, 13] There is an abundance of literature on structure, function and molecular biology of Ras proteins.[14] Multiple efforts have focused on the development of affinity binders to target K-Ras proteins.[15, 16] However, K-Ras has proved to be a difficult target due to its intracellular localization, presence of multiple isoforms, and lack of druggable pockets.[17] Recently, Amgen [18] and Mirati [19] therapeutics have developed covalent Ras inhibitors to the G12C mutant due to the reactivity of the cysteine mutation as a nucleophile. But this strategy cannot be extended to the other mutations, and there’s a still need for developing affinity molecules to target Ras proteins.

In this work, we developed VHH binders against the most common K-Ras mutant (G12D) from a camelid derived VHH-library using phage display and identified lead hits that showed binding on phage ELISA. However, we found that most of the lead hits showed poor soluble expression in bacterial cultures, which limited their characterization and functional studies. We hypothesized that the poor solubility is due to the high surface hydrophobicity of the clones, which can be improved by hydrophilic surface mutations. To that end, we employed a combination of random and in silico guided mutagenesis. Specifically, the in-silico approach involved a computational pipeline for modeling the 3D structure of the lead hits and identifying hydrophobic hotspots on the surface of the binders. By using spatial aggregation propensity (SAP) as a measure for surface hydrophobicity, we showed that the clones with poor soluble expression contained significant hydrophobic hotspots.

Having shown correlation between the hydrophobicity and expression levels, we followed two approaches to perform mutations and produce high expression clones. i) mutating hydrophobic amino acids centered at the hydrophobic hotspots to charged or polar ones; ii) combining mutations that are identified through a round of random mutagenesis, which are predicted to improve surface hydrophobicity. Both approaches led to significant increase in soluble expression (2x to 40x) of the binders, and therefore allowed characterization of their affinities to their target. While most of the binders produced by the former approach suffered from loss of binding affinity, the latter approach resulted in high solubility without loss of binding affinity. Therefore, based on our results, we recommend performing a round of random mutagenesis to identify mutations, and using an *in-silico* pipeline to guide the evolution of the binders by simulating several combinations of the mutations and selecting for soluble expression. Overall, our work provides a framework for rational design to improve expression of VHH binders.

## 2. Materials and methods

### 2.1 Homology modeling and all-atom MD simulations

Modeling and simulations were performed for predictions of surface hydrophobicity of the VHH binders. The VHH binders were modeled using homology modeling, followed by energy minimization and equilibration in all-atom MD simulations in water. The Molecular Operating Environment (MOE) [20, 21] and Iterative Threading ASSEmbly Refinement (I-TASSER 5.1) [22] software were used for the homology modeling, and NAMD parallel-computing code (NAMD 2.12) [23] and GROningen MAchine for Computer Simulations (GROMACS-2018) [24] were used for all-atom simulations. A 5 ns equilibration run was followed by a 100 ns of production run. The MD simulations incorporated leap-frog algorithm with a 2 fs time-step to integrate the equations of motion. The system was maintained at 300K and 1 bar, using the velocity rescaling thermostat [25] and Parrinello-Rahman barostat [26], respectively. The long-ranged electrostatic interactions were calculated using particle mesh Ewald (PME) [27] algorithm with a real space cut-off of 1.2 nm. LJ interactions were also truncated at 1.2 nm. TIP3P model [28] was used represent the water molecules, and LINCS [29] algorithm was used to constrain the motion of hydrogen atoms bonded to heavy atoms. Co-ordinates of the protein molecule were stored every 1 ps for further analysis.

### 2.2 SAP calculations

Following a similar protocol as Trout and coworkers [30], we used all-atom simulations of the VHH binders in explicit water, and calculated SAP averaged over the coordinates obtained in a 100 ns simulation trajectory, as follows

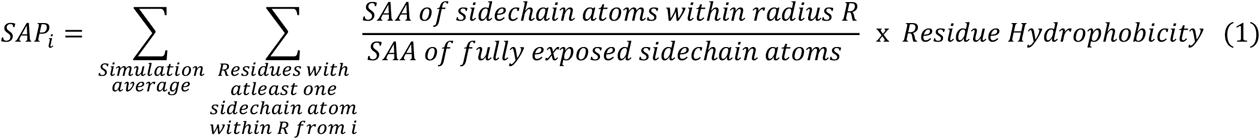

The equation (1) allows calculation of residue specific spatial aggregation propensity (SAP_i_) based on normalized solvent accessible areas (SAA) of the side chain atoms in the vicinity (defined by a distance cut-off, R) and inherent residue hydrophobicity (from Kyte-Doolittle hydropathy scale).[31]

### 2.3 VHH – periplasmic expression

pADL-VHH plasmids were transformed into competent *E. coli* BL21 (DE3) cells and grown on Luria Bertani (LB) Agar plates. Single colonies were selected using pipette tips and grown in 10 mL of LB + Glc + Carbenicillin starter culture overnight at 37 °C. For VHH protein expression the starter cultures were diluted in 500 mL Terrific Broth (TB), and grown at 37°C, followed by the induction with 1 mM IPTG once the O.D_600_ reached 0.6 and incubated at 30°C overnight. This was followed by an osmotic shock to selectively release the periplasmic proteins into the supernatant, which was stabilized by high sucrose concentration. The periplasmic fraction is harvested by centrifugation and the recombinant His-tagged proteins were successively purified through immobilized metal affinity chromatography (NiNTA resin) and size exclusion column via Fast Protein Liquid chromatography (FPLC, Superose 6 Increase 10/300 GL column) Briefly, the harvested fraction was incubated with the nickel activated affinity resin (NiNTA) and following several washes, the His-tagged proteins were finally eluted in a highly-concentrated imidazole buffer and stored at 4°C. The presence of VHH was assessed on a 4-12 % Bis-Tris gel. VHH proteins were concentrated, and buffer exchanged in PBS before a final FPLC purification step (ÄKTA pure instrument; GE Healthcare Life Sciences) via a small-scale size exclusion chromatography column (Superdex 200 Increase 10/300 GL; GE Healthcare Life Sciences). The protein of interest was further eluted via isocratic elution in PBS buffer containing; 25mM trehalose, 0.5mM MgCl2, 0.05% Tween20before being concentrated, aliquoted and stored at -80C. The final concentration of the protein was determined using 280nm absorbance on Nanodrop and using storage buffer as a blank. The protein concentration was then adjusted to perform characterization assays.

### 2.4 Enzyme Linked Immuno Sorbent Assay (ELISA)

The VHHs in their soluble form were characterized for their selective binding using ELISA performed on 96 well plates. The wells were coated overnight with K-Ras G12D antigens or PBS (for negative control). The coated wells were then washed with PBS and 0.05% Tween four times and blocked with 2% Milk-PBS. The VHHs were introduced in the wells serially diluted in blocking buffer in the range from 24.4 ng/ml (∼1.5nM) to 100 ug/mL (∼6.25nM) including a blank with no protein for further background subtraction. Anti-Ras antibody (Abcam, ab55391) was used as positive controls in a concentration range starting from 0 to 2ug/mL. Following incubation of the targeted antigen (just blocking buffer) with the variable VHH concentrations, the plate was washed with PBS, 0.05% Tween to remove unbound VHH. To detect the antigens bound VHH, an anti-6x His tag primary Mab (Thermo Fisher Scientific 4E3D10H2/E3) was introduced in 1:1000 dilution and incubated for one hour before further washes. Finally, a goat anti-mouse IgG polyclonal secondary antibody (Invitrogen) was finally added in 1:5000 dilution and incubated at room temperature before signal development. After final washing steps, horse radish peroxidase substrate was added and the absorbance at 450 nm was read on Tecan 20M Spark plate reader following stoppage of the reaction with 0.16M H_2_SO_4_. Each condition was performed in duplicate or triplicate and absorbance signals were analyzed based on the average signal calculated on two or three wells. Standard deviation was determined based on duplicated or triplicated conditions as well.

### 2.5 Bio-layer Interferometry (BLI)/Affinity Determination

Kinetic assays were carried out at 30°C with orbital shaking at 1,000 rpm on a ForteBio Octet RED384 instrument (Molecular Devices) with a running buffer of 1X Kinetics buffer (Sartorius) + 0.25% BSA and using ForteBio Data Acquisition software 9.0. Random Amine chemistry with Amine Coupling Second Generation kit (Sartorius) was used to load K-Ras G12D ligands on Amine Reactive Second Generation (AR2G) biosensors (Sartorius) according to the manufacturer’s instructions. Briefly AR2G sensors equilibrated in water for 10 minutes, were activated with 1-Ethyl-3-[3-dimethylaminopropyl] carbodiimide hydrochloride) (EDC) and s-NHS (N-hydroxysulfosuccinimide) (NHS) for 300 seconds and were either mock loaded (no ligand reference sensor) or loaded with 30ug/ml of K-Ras G12D in sodium acetate buffer, pH 5.0 for 600 seconds followed by a quench step with 1M ethanolamine pH 8.5 for 500 sec. Each ligand loaded sensor equilibrated in running buffer to establish a baseline, were dipped into wells containing 2-fold dilutions of VHH analytes (F4.0, F4.3, F4.6, B6.0 and B6.5) in running buffer, or buffer alone (reference sample). Increasing concentrations of analytes ranging from 15µM-0µM were allowed to associate for 300 secs, followed by a 600 sec dissociation step, where the VHH-bound sensors were washed with running buffer. Binding curves were analyzed using ForteBio Data Analysis HT 10.0 evaluation software. To control for background signal and non-specific binding, raw experimental data were processed by subtracting reference biosensor (no ligand) and reference sample (no analyte). Sensors were regenerated with two short 15 seconds pulses of 10 mM Glycine pH 1.7 followed by 180 seconds of wash with running buffer. For each interaction pair, processed data from three independent experiments of at least five different concentrations of analyte binding were globally fit to a 1:1 Langmuir binding model (or heterogenous ligand binding model) to calculate the association rate constant *k*_a_, the dissociation rate constant *k*_d_. Equilibrium dissociation constants *k*_D_ which defines the strength of the interaction or affinity was calculated as the kinetic dissociation rate constant divided by the kinetic association rate constant.

## 3. Results

Our primary goal was to develop VHH binders against K-Ras G12D mutant. Using phage display, we performed a screen on a camelid derived naïve VHH library and selected the clones that showed preferential binding to the target as lead hits. The sequences of the four lead clones (F4.0, F3.0, B6.0 and E4.0) and their alignments shown in **Fig. 1A**, confirmed uniqueness of the clones as highlighted by substantial differences in the CDR regions. We proceeded to express these clones in bacterial expression cell cultures (see Methods section), and found very poor yields, as shown in **Fig. 1B**. Traditionally, VHHs are thought to be stable and highly soluble, and hence are expected to possess high levels of expression. However, in repeated expression of our lead hits, we found very minimum yields. We hypothesized that the naïve library created by combinatorial methods potentially resulted in the low expressing clones, as the surface hydrophobicity is not controlled for in their generation.

**Figure 1:**
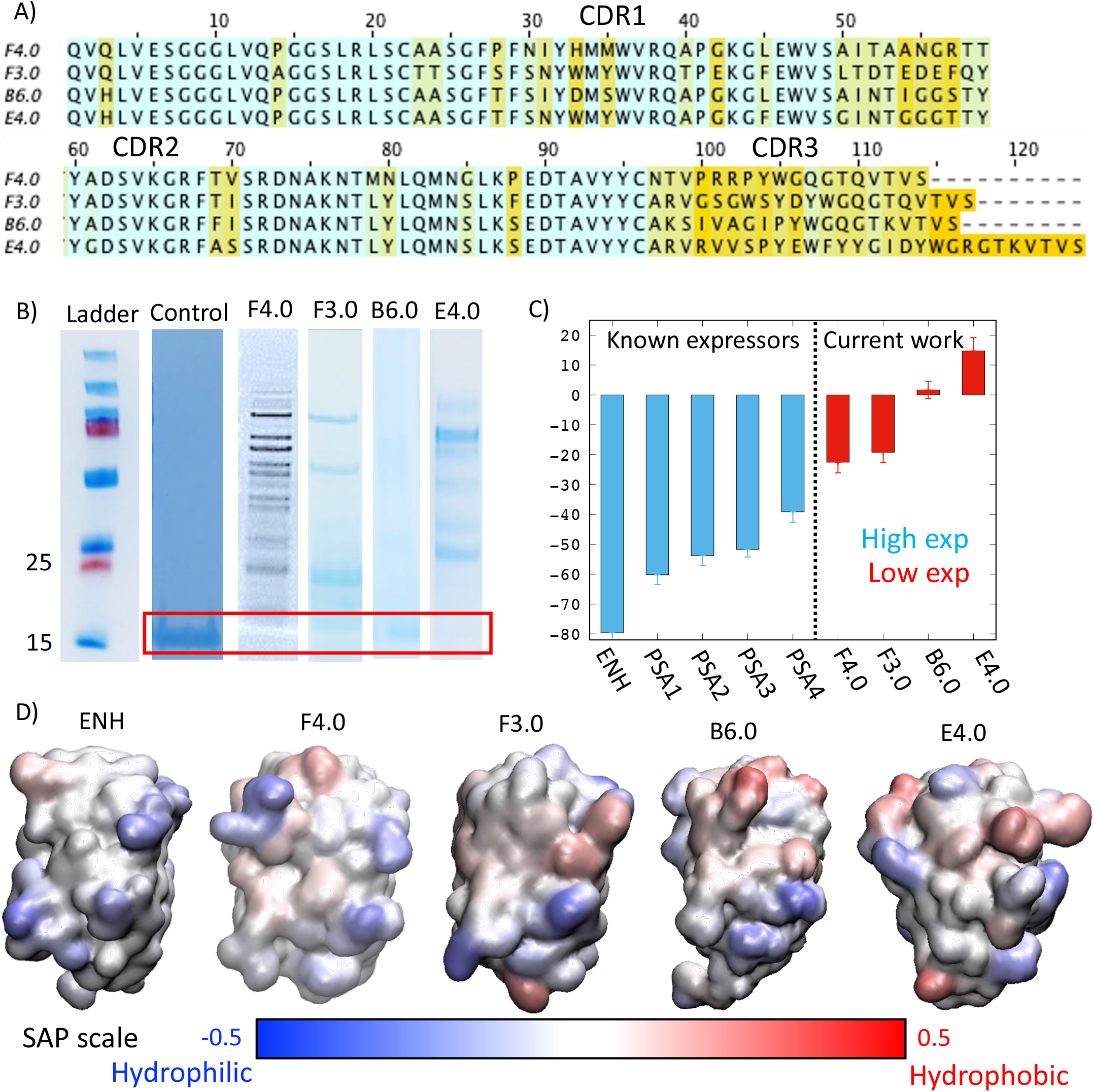
A) Sequence alignment of the lead clones identified from Phage ELISA B) Gels showing poor soluble expression of the VHH clones C) Time-averaged SAP scores for a range of VHH binders from literature to the ones generated in this work. D) SAP maps for the VHHs delineating hydrophobic and hydrophilic regions of the surface. The side chosen exposes the CDRs and the other face shows little variability and hence not shown.

To test the above-mentioned hypothesis, we adopted a computational approach developed by Trout and co-workers for improving stability of antibody binders. In their work, Trout and co-workers [30] developed a molecular context dependent measure for surface hydrophobicity of proteins, called Spatial Aggregation Propensity (SAP), which can be characterized using MD simulation trajectories. The 3D structures of VHHs needed for this approach was obtained using a combination of MOE and I-TASSER for consensus, which was then subject to energy minimization, equilibration and a 100 ns production run in explicit water to arrive at an ensemble of representative structures for SAP calculations. The SAP_*i*_ score for i^th^ structure was calculated using **Equation 1**, which was then used for obtaining i) the overall SAP score, a summation obtained over all the residues of the VHH (**Fig. 1C**), and ii) SAP maps that identify hydrophobic hotspots, highlighting the locations that can be targeted for solubility enhancing mutations (**Fig. 1D**).

In **Fig. 1C**, the SAP score of a GFP-specific VHH, Enhancer (ENH) and a few other known VHH binders to prostate specific antigen (PSA) were compared with the K-Ras specific lead hits from our phage display screen. Interestingly, ENH, which is commercially available, and a known high expressing protein showed significantly lower SAP score compared to our clones. In addition to ENH, we also observed significantly lower SAP scores for the high expressing PSA-specific VHHs, indicating a correlation between SAP scores and soluble expression. This observation suggested likelihood of hydrophobic patches on the surface of the K-Ras specific binders from our screen, which might lead to aggregation and consequently lower expression of the proteins. The SAP maps for the VHH binders shown in **Fig. 1D** supports this hypothesis, highlighting the hydrophobic patches on low-expressing clones, especially for clones B6.0 and E4.0. These maps also provide guidance for targeted mutagenesis of the binders for soluble expression.

To improve the soluble expression of the binders, we followed two different approaches with two select clones (B6.0 and F4.0, respectively). Our first approach is purely computational and involved introducing targeted hydrophilic (charged/polar) mutations in the hydrophobic hotspots of B6.0 while minimizing the impact to the CDR regions. Specifically, we identified six positions in the B6.0 sequence: I31, I54, Y59, F69, Y106 and Y107, that form hydrophobic patches on the surface of the protein as identified in **Fig. 2C**. The sequences of the mutated clones (B6.1 to B6.6) are shown in **Fig. 2A**, containing two to six mutations, with varying levels of hydrophobicity. The soluble expressions and the SAP scores are shown in **Fig. 2B** and **Fig. 2D**, respectively. While two of the clones (B6.1 and B6.4) did not show any improvement in soluble expression, the other four clones (B6.2, B6.3, B6.5 and B6.6) had remarkable increase in yields (2x to 8x) and were expressed at high enough levels for further characterization. This is remarkable considering the high expression was achieved with just a few select of mutations. Interestingly, we were also able to establish a strong correlation (R^2^=0.98) between the SAP scores and the soluble expression of the VHH clones (**Fig. 2E**), except for clone B6.4, which implies there might be additional factors at play, such as protein folding. Nevertheless, by introducing a few hydrophilic mutations at the hydrophobic hotspots, we were able to significantly increase the expression of the VHH. Past theoretical studies focused on surface hydrophobicity at the nanoscale have predicted an asymmetric effect of large hydrophobicity changes from small hydrophilic perturbations, which are consistent with our observations here.[32]

**Figure 2:**
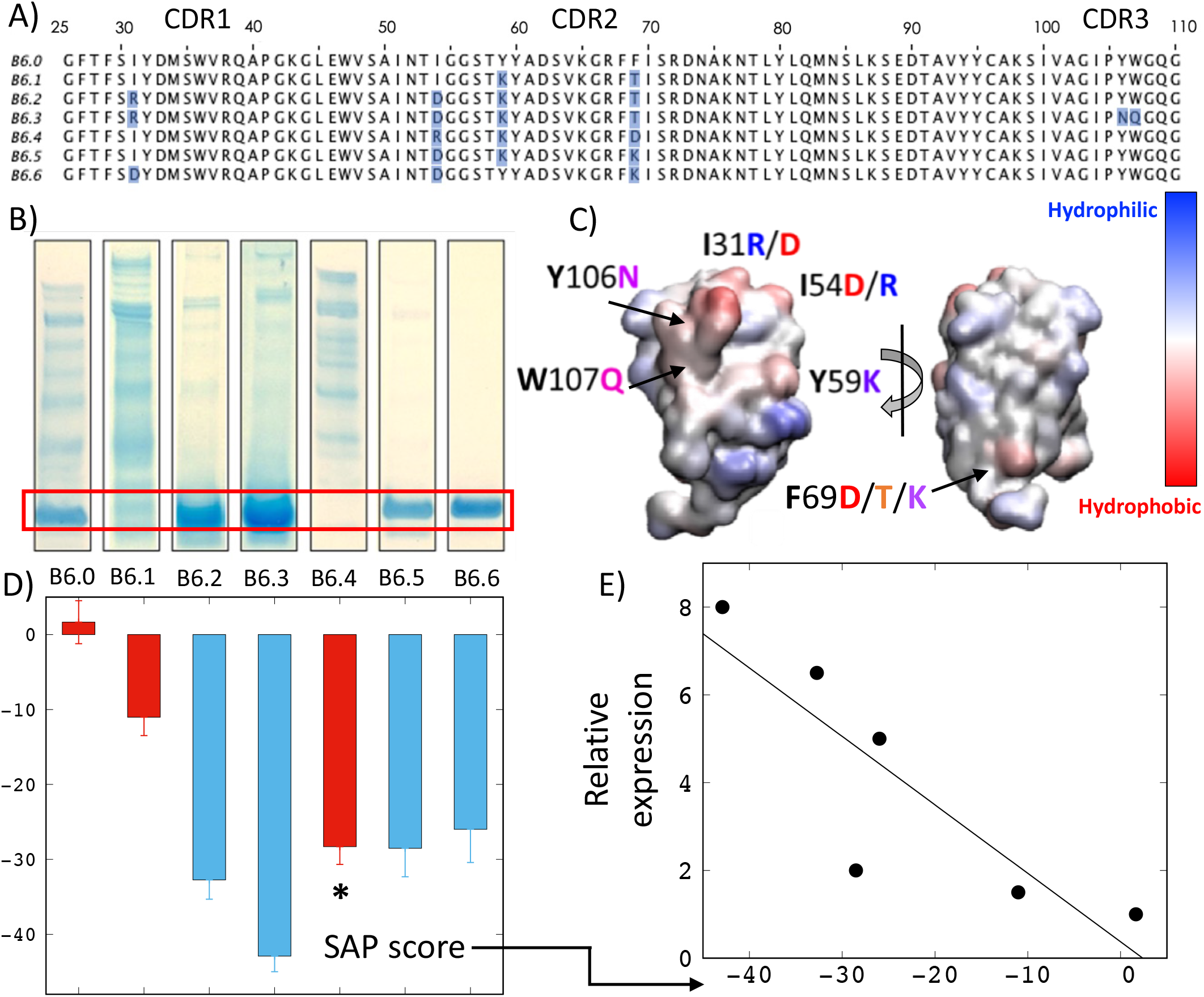
A) Amino acid sequences alignment of B6 clones with mutations highlighted. B) SAP maps of B6 clones. The residues of interest are marked, and the hydrophilic point mutations are highlighted. C) Expression and D) SAP scores of the B6 clones. E) The plot of relative expression (C) versus SAP scores (D) showing a strong correlation.

Alternatively, following a different approach, we performed one round of random mutagenesis on the F4.0 using phage display, and identified three clones (F4.1, F4.2 and F4.3) that either improved or maintained the affinity to the target (K-Ras). While clones, F4.1 and F4.2 had single mutations, I55N and G109D, F4.3 had two mutations: W71R and R125P (**Fig. 3A** and **Fig. 3B**). We performed expression of the new clones and found a nominal increase in the expression level for the F4.2 and F4.3 clones (**Fig. 3C**). We modeled these clones and found that the SAP scores improved significantly for the best expressing clone (F4.3). Interestingly, this improvement is facilitated by a single amino acid hydrophilic mutation (W71R) in the middle of a hydrophobic patch as highlighted in **Fig. 3B** (F4.0 vs F4.3). We modeled three novel clones F4.4, F4.5 and F4.6 (sequences shown in **Fig. 3A**) that combined the mutations from F4.1, F4.2 and F4.3. We found significant reduction in SAP scores for the novel clones, with F4.6 showing the largest decrease in surface hydrophobicity. We expressed of all the novel clones and found all of them had significant increases in expression levels, consistent with the SAP scores. Remarkably, F4.6 had a 40x increase compared to F4.3 (**Fig. 3E**). The surface maps of the clones colored by SAP scores clearly show elimination of the hydrophobic patches by addition of hydrophilic side chain amino acids. Due to the increased levels of expression from both these approaches, we were able to further characterize these clones on soluble ELISA and Octet.

**Figure 3:**
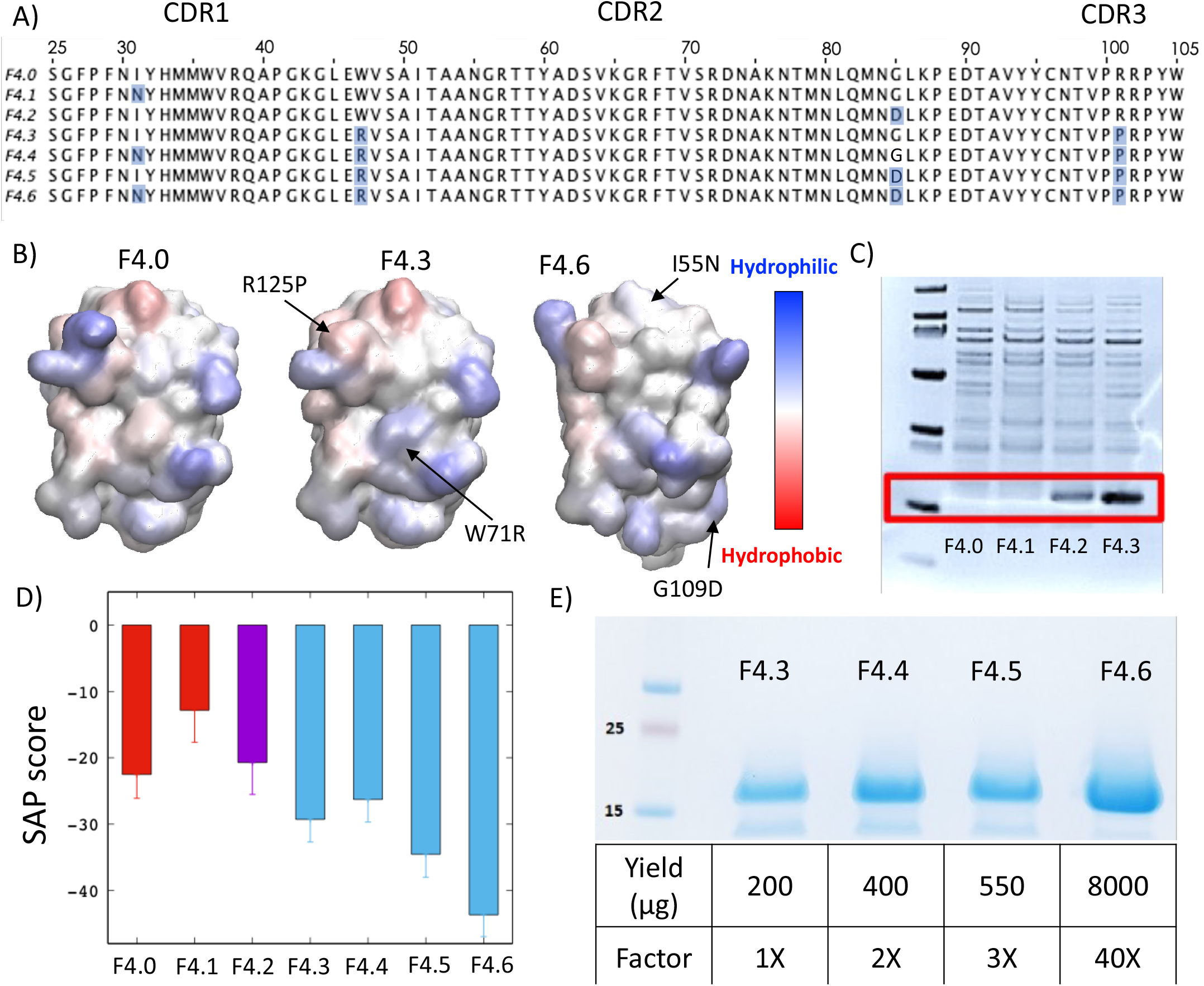
A) Amino acid sequences alignment of F4 clones with mutations highlighted. B) SAP maps of few select F4 clones with hydrophobic regions highlighted in red. C) Expression profiles of the first set of F4 clones. D) SAP scores of all F4 clones. E) Expression profiles and quantification of yield for the improved F4 clones.

The moderate to high expressing clones (five B6 and four F4 clones) were produced in 500 ml cultures. First, we performed Enzyme Linked ImmunoSorbent Assay (ELISA) of the clones in their soluble form against the target (**Fig. 4**). We adsorbed K-Ras G12D on to a surface and incubated them with VHHs overnight. His tags on the VHHs which were added for purification, were also used in their recognition with anti-His antibodies. While all the F4 clones showed similar affinity towards the target in the sub μM range, most of the B6 clones (except for B6.5) showed decreased binding (**Fig. 4B** and **Fig. 4C**). This behavior clearly highlights the shortcomings of our first approach of performing hydrophilic mutations and indicates the need for random mutagenesis to identify potential soluble enhancing mutations. We chose B6.0, B6.5, F4.3 and F4.6 for further analysis to quantify the binding affinity.

**Figure 4:**
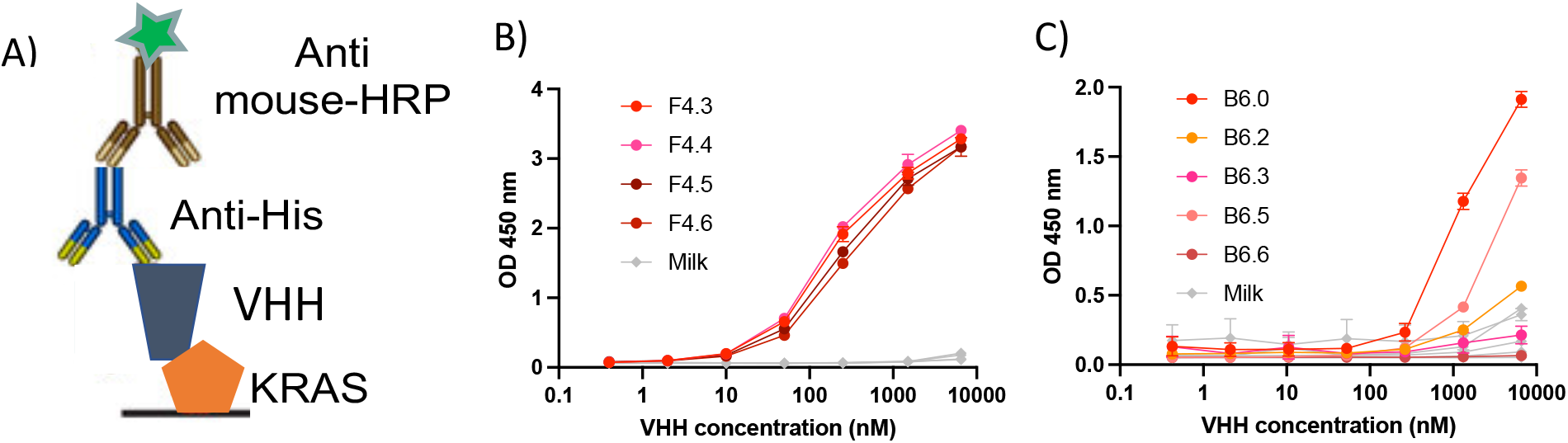
A) Schematic representing standard ELISA for characterizing VHH binding to K-Ras. B) and C) ELISA curves of F4 and B6 clones, respectively.

To quantify the strength of the interactions between the VHHs and K-Ras G12D, we performed binding assays on the Octet instrument and quantified the affinity of select clones to the target. Briefly, we loaded K-Ras G12D on the sensor tip using random amine adsorption to achieve a varied number of orientations of the antigen on the surface. The VHH was then introduced at various concentrations, and binding of the VHH to the surface is then monitored over 300 seconds, followed by a buffer wash over 600 seconds, where the unbinding kinetics is monitored. The curves thus obtained at various concentrations were fit to Langmuir isotherms using a 1:1 binding model, to arrive at rate constants and the equilibrium dissociation constant, k_D_ shown in **Table 1**.

**Table 1:** Kinetics and thermodynamics of various VHHs binding to K-Ras at 30°C.

| VHH | $k_a$ (Ms)<sup>-1</sup> | $k_d$ (s<sup>-1</sup>) | $k_D$ (nM) |
| --- | --- | --- | --- |
| B6.0 | $(28.35 \pm 0.65) \times 10^3$ | $(46.00 \pm 0.34) \times 10^{-3}$ | $1730 \pm 39$ |
| B6.5 | $(9.24 \pm 0.34) \times 10^3$ | $(189.0 \pm 2.35) \times 10^{-3}$ | $20500 \pm 804$ |
| F4.3 | $(48.44 \pm 1.43) \times 10^3$ | $(9.71 \pm 0.06) \times 10^{-3}$ | $199 \pm 6.0$ |
| F4.6 | $(32.31 \pm 0.53) \times 10^3$ | $(3.11 \pm 0.01) \times 10^{-3}$ | $96 \pm 1.6$ |

Kinetic rate constants were determined using 1:1 Langmuir binding model in ForteBio Data Analysis HT 10.0 evaluation software. $k_a$ is the association rate constant, $k_d$ is the dissociation rate constant, $k_D$ is the equilibration dissociation constant. Each experiment was carried out in triplicates with at least five different concentrations of the VHH ranging from 0.4-15 μM.

Consistent with the ELISA results, we saw stronger binder affinity (as indicated by k_D_) for the F4 clones compared to the B6 clones. While B6.5 showed a μM range affinity to the target, the F4.6 clone possessed an impressive sub 100 nM affinity, which is an improvement from its parent clone (F4.3). The kinetic constants reveal weaker association of both B6.5 and F4.6 compared to their respective parent clones as expected from the hydrophilic nature of the mutations, which might increase the tendency of the protein to remain in an unbound state. However, while the dissociation kinetic constant is also weaker for the B6.5 clone, the F4.6 clone gains significant interactions with the target and remains strongly bound. Therefore, the high expressing F4 clone (F4.6) holds potential to be a K-Ras binder, especially in a multimeric form. We are currently pursuing to produce affinity reagents for labelling K-Ras G12D in cellular imaging.

## 4. Discussion and Conclusion

In this work, we have tested a novel computationally guided approach to improve soluble expression of VHH binders. Our work is motivated by poor expression of VHH binders selected from a naïve phage library screened against our target (K-Ras). Similar observations have been made by other researchers, who have proposed ways to improve stability and expression of VHH binders.[33] We hypothesized that the high surface hydrophobicity of the VHHs screened in our study contributes towards poor solubility, leading to poor soluble expression. To support this hypothesis, we adapted earlier work by Trout and coworkers,[30] in which they developed a computational molecular measure of surface hydrophobicity called Spatial Aggregation Propensity (SAP). Through modeling and simulations, we were able to produce SAP maps and average SAP score, which not only showed high SAP scores for poorly expressing clones but also established a strong correlation of surface hydrophobicity with levels of soluble expression. To our knowledge, this is the first study to establish a correlation between SAP scores and expression levels of VHH binders.

Having established this correlation, we set out to improve soluble expression of these binders that suffer from poor expression. Based on previous studies of hydrophobicity at the molecular scale, hydrophobic hotspots can be neutralized by single amino acid mutations.[32] Therefore, we hypothesized introducing a few surface mutations will result in significantly lower SAP scores. We followed two different approaches to perform mutations. In the first approach, we tested a few hydrophilic mutations that are predicted to improve SAP scores. This approach is purely computational, and the mutations were arbitrarily chosen to be charged/polar. As we expected, most of these clones had higher expression, showing a strong correlation between SAP scores and expression levels. While this approach showed promise, most of the high expressing clones also lost their binding affinity to their target when they were tested on ELISA. Even though the mutations sites were carefully chosen not to be in the CDR regions, we suspect that due to the small size of the VHH, the regions proximal to the CDR regions are also involved in its binding to the target. Even for the B6.5 clone, which had comparable binding on ELISA, the equilibration dissociation constant was significantly higher than the parent clone (B6.0), indicating loss of binding affinity.

In an alternate approach, we performed a round of random mutagenesis on phage display and selected a few clones (with one or two mutations) that showed comparable binding to the target. Some of these clones showed marginal increase in expression levels and provided us with a list of mutations that can be combined computationally and tested. In doing so, we predicted that the clone that combined all the four mutations would have the highest expression level. Remarkably, the soluble expression was completely in agreement with the predictions. Importantly, these clones showed no loss in binding affinity to the target on ELISA, and even showed stronger binding indicated by the lower equilibration dissociation constant.

In conclusion, we have presented a hybrid approach that combined random mutagenesis with computational predictions to improve the soluble expression of VHH binders. Even though similar studies have been conducted on antibodies (IgG),[30] this is the first application of such an approach VHH binders. Our study shows that VHH binders need this hybrid approach to arrive at solubility enhancing mutations due to their smaller size compared to IgGs. Even mutations in the non CDR regions, while improving expression levels, might lead to loss of binding and hence need to be chosen carefully through random mutagenesis.

## Acknowledgements

This work was supported by the Cancer Early Detection Advanced Research Center at Oregon Health & Science University (Full6130619). All simulations had been performed using the high-performance computational facilities an Exacloud cluster at Oregon Health & Science University (Portland, OR).

